# Linking Demyelination to Compound Action Potential Dispersion with a Spike-Diffuse-Spike Approach

**DOI:** 10.1101/501379

**Authors:** Richard Naud, André Longtin

## Abstract

To establish and exploit novel biomarkers of demyelinating diseases requires a mechanistic understanding of axonal propagation. Here, we present a novel computational framework called the stochastic spike-diffuse-spike (SSDS) model for assessing the effects of demyelination on axonal transmission. It models transmission through nodal and internodal compartments with two types of operations: a stochastic integrate-and-fire operation captures nodal excitability and a linear filtering operation describes internodal propagation. The effects of demyelinated segments on the probability of transmission, transmission delay and spike time jitter are explored. We argue that demyelination induced impedance mismatch prevents propagation mostly when the action potential leaves a demyelinated region, not when it enters a demyelinated region. In addition, we model sodium channel remodeling as a homeostatic control of nodal excitability. We find that the effects of mild demyelination on transmission probability and delay can be largely counterbalanced by an increase in excitability at the nodes surrounding the demyelination. The spike timing jitter, however, reflects the level of demyelination whether excitability is fixed or is allowed to change in compensation. This jitter can accumulate over long axons and leads to a broadening of the compound action potential, linking microscopic defects to a mesoscopic observable. Our findings articulate why action potential jitter and compound action potential dispersion can serve as potential markers of weak and sporadic demyelination.

## Introduction

Axons are the only link between our brains and the external world. Whether through touch, smell or vision, all the information we have about the external world is channeled through bundles of axons. Similarly, our only mean to interact with the external world is to send muscle commands along another set of axons. Clearly, this link must be made as fast as possible and myelination has evolved to regulate axonal propagation, minimizing the ‘*temps perdu*’ as coined by Hermann von Helmholtz [1]. Given its essential role in brain function, it is perhaps not surprising that disorders of axonal myelination are linked to dramatic loss of function. It is not clear, however, how mild demyelination affects propagation along nerve bundles, since its effect on internodal transmission is subtle. To better understand how mild demyelination gives rise to the early symptoms of demyelinating diseases, one approach is to build mathematical models of the axon and study the link between demyelination patterns and axonal conduction properties [2, 3, 4, 5].

Mathematical modelling of myelinated axons can be separated into two approaches. The first approach, pioneered by Fitzhugh (1961) [6], aims to determine the basic electrodynamic properties regulating the flow of charges along a myelinated membrane. The propagation dynamics between the nodes are encapsulated in a single equation called the cable equation, which can be solved either analytically or with numerical methods to obtain attenuation or filtering properties [7, 8]. This approach is restricted to uniform segments; it cannot typically capture the inhomogeneous myelination, paranodes or the alternation with spiking nodes of Ranvier. To capture these inhomogeneities, one uses a second approach called compartmental modeling, where the system is modeled using a large number of small, locally uniform segments. Simulating numerically the exchange of charges between segments with different dynamic properties has revealed the important roles of myelin morphology and nodal dynamics for the propagation of action potentials [9, 10, 11, 4, 12]. This has been the main method to address effects of demyelination [2, 3, 5], ion channel noise [13, 14, 15, 16], and spike after-potential [4] on axonal propagation.

In the present article we consider a hybrid approach. We model sequentially the nodal action potential generation in a small locally uniform spiking compartment and the internodal propagation resulting from a cable equation. Inspired from recent work on stochastic-integrate-and-fire models [17, 18, 19, 20, 21, 22, 23, 24] and the spike-diffuse-spike model of dendritic spine activation [25], we develop here the stochastic spike-diffuse-spike model (SSDS). We use this framework to investigate the propagation along axons where the excitability of a node can be enhanced to compensate for demyelination. We will start by describing the modelling framework and how it incorporates demyelination as well as homeostatic ion channel re-insertion. The framework will then be simulated to estimate the effect of mild and sporadic demyelination on the features of propagation. We will compute delay and jitter of action potential propagation first across a single node and then along a long axon. In addition, we will compute these metrics in the presence of homeostatic regulation of nodal excitability. Finally, we will use these estimates to compute the width of a compound action potential in various demyelination conditions. Potentially an early marker of demyelinating diseases [26, 27, 28], our results articulate how weak and sporadic axonal damage affect the delay and dispersion of the compound action potential.

## Methods

The goal of this section is to expose the computational framework. First, we describe SSDS and use it to calculate analytically multiple quantities of interest: conduction speed, conduction probability and spike timing jitter. Once the basic propagation model is layed out, a model of demyelinating damage and articulate how this damage influences conduction speed, probability and jitter. Subsequently, the properties derived herein form the basis of the propagation along an axon bundle. We relate the single axon model to the characteristics of the compound action potential last.

### Single Axon Model: Stochastic Spike-Diffuse-Spike

We consider axons as an alternation of Ranvier nodes and myelinated segments. The Ranvier node is strongly excitable but very small. In contrast, the internodal region is not excitable, but extends spatially. For that reason, Ranvier nodes are modelled by an excitable compartment with no spatial extent and the internodal segment is modelled by a passive compartment with spatial extent. This modelling framework has been introduced previously for active dendritic spines scattered along a dendrite [25]. Here we extend this modelling framework to capture axonal properties with the use of kernel-based methods. The two steps are described mathematically as

1. **Internodal Drift-Diffusion**. Given an action potential leaving a node (Fig. 1 (a)), the charges entering and leaving the axon at the node will propagate along the myelinated internodal region to the next Ranvier node. This operation is called *drift-diffusion* because a large and succinct current in one node will be felt as a weaker and longer-lasting depolarization at the next Ranvier node, similar to the time evolution of molecules in water in the presence of a drift. At the location of the next Ranvier node, the result of this operation is depolariztion of the membrane potential (Fig. 1 (b)).
2. **Nodal Spiking**. Given the membrane potential depolarization, the next Ranvier node will fire an action potential with probability proportional to the distance between this depolarization and a membrane potential threshold (Fig. 1 (c)).

**Figure 1.**
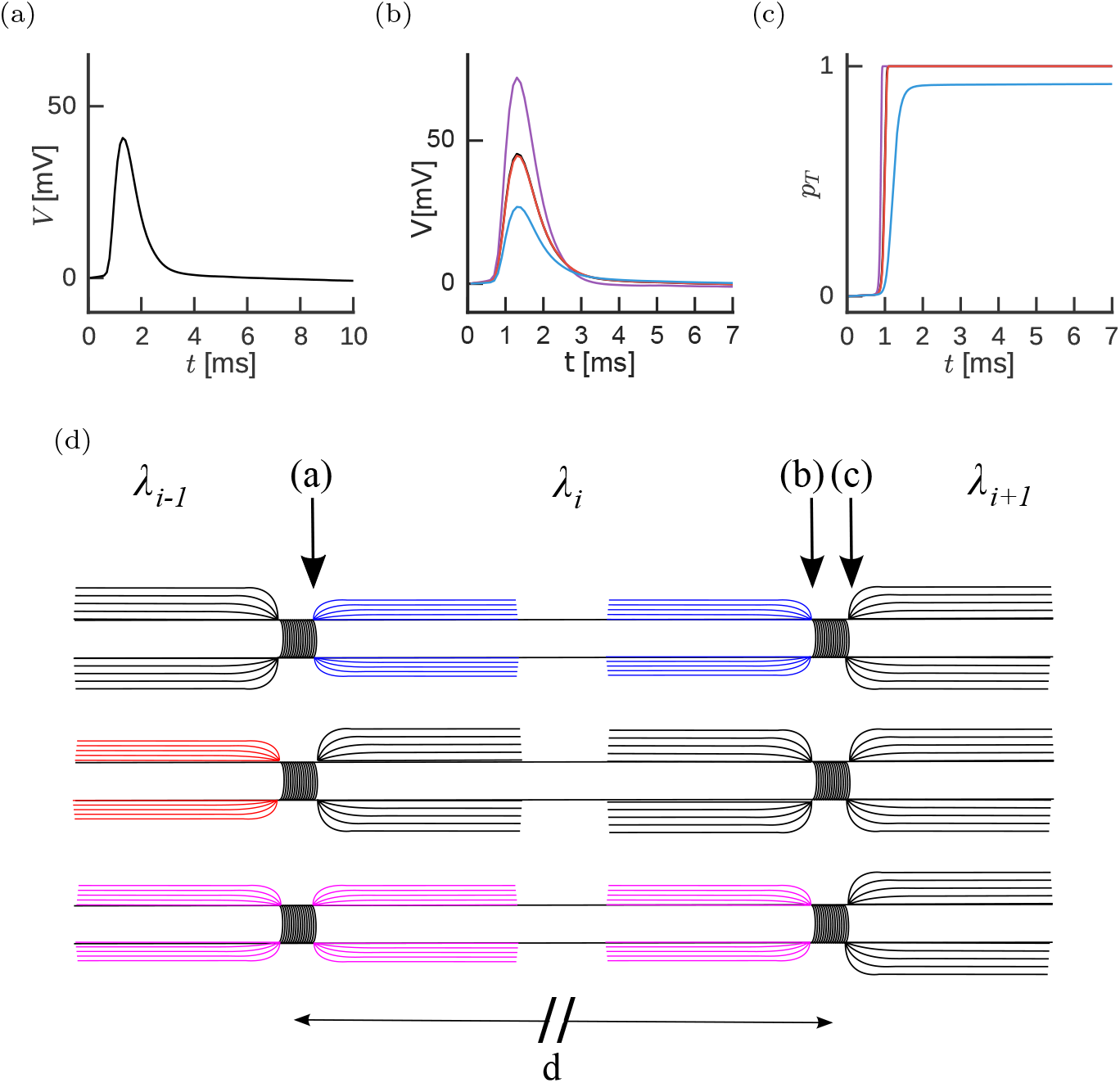
The Stochastic Spike-Diffuse-Spike Model. (a) The action potential shape as it leaves the Ranvier node, schematically indicated in (d). (b) Propagation through the next internode alters the action potential shape as it reaches the next node (black, almost confounded with red). Damage to the internodes will affect this propagation following one of three possibilities. We consider whether demyelination affects the orthodromic internode only (purple), the antidromic internode only (blue) or both internodes equally (red). (c) Probability of transmission as a function of time for the intact and the three cases of damage. (d) Schematic illustration of two Ranvier nodes separated by a distance d for the three damage configurations. For a propagation from left to right, we consider three possibilities for an action potential starting at node (a). Top: demyelination of orthodromic internode. Middle: demyelination of antidromic internode. Bottom: equal demyelination of both anti- and orthodromic internodes.

We alternate steps (1) and (2) to reproduce the process of saltatory conduction. In the following, we describe each of these steps in detail, along with the rationale for our modeling choices. For clarity of exposition, our simplifying assumptions are summarized here:

1. **Action potential shape is invariant**. Widening and shortening of action potentials produced at high frequency is therefore not captured by our present model.
2. **Axons are modeled as a single thread**. Reflection and impedance mismatch at bifurcations would require extensions to the present model.
3. **The electric coupling between nodes is restricted to pairs**. For real axons, however, nodal spiking has a small but systematic effect multiple internodes away.
4. **Propagation of current along an internode follows an idealized, semi-infinite, uniform cable**. Taking into account the impact of peri-nodal structure and other types of inhomogeneous myelination would require a modification of the model. We have assumed a lumped compartment at the origin, composed of a Ranvier node and the antidromic compartment assumed to remain isopotential.
5. **Demyelination can be modeled by changes in the electrotonic length constant**. The electrotonic length constant is the distance over which a depolarization has decayed by a factor of *e*^−1^, at steady state. Myelin greatly increases this length constant. The effects of demyelination on other parameters-such as the membrane time constant-are assumed to play a much weaker role [29].

#### Internodal Drift Diffusion

We want to relate the current associated with a spike *leaving* a node *I*_AP_(*t*) (Fig. 1 (a)) to the membrane potential that this action potential causes in the *next* Ranvier node (Fig. 1 (b)). The effect of charged molecules drift-diffusing along the cable with potential leak across the membrane is captured by a convolution of *I*_AP_(*t*) with the Green’s function, or impulse-response function of the internodal region. The Green’s function takes into account the dynamics of the cable as well as possible impedance mismatch of the paranodal regions. Although the Green’s function is typically defined for all points in space, we are interested in the membrane potential at the location of the next node. Assuming that the orthodromic position from node *i* can be parametrized by a single coordinate *X*, we let *G_i_*(*T, X*) be the Green’s function of the ith internodal region. Accordingly, we can compute the membrane potential along the ith internode *V_i_*(*T, X*) with origin at the ith Ranvier node as

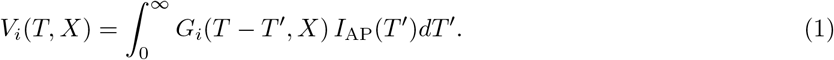

where *X* = *x*/λ*_i_* is in units of the electrotonic length of the orthodromic internode λ*_i_* and *T* = *t*/*τ* in units of the membrane time constant *τ*.

Analytical expressions of *G* have been derived for a cable that is either uniform passive [21], non-uniform [30] or quasi-active [31]. Otherwise, the detailed geometry of the paranodes can be taken into account by computing the impulse-response function numerically [32]. Alternatively, one could use an empirical estimate of the impulse response function. For clarity of the exposition, we will limit the present analysis to an analytical expression of the Green’s function corresponding to a uniform, semi-infinite and passive cable with a compartment lumped at the origin of the cable. This expression was derived for propagation of charges in a uniform semi-infinite and passive dendrite with a somatic compartment lumped at its base [33]. The assumptions underlying this derivation are also satisfied by the axon leaving a leakier Ranvier node. The Green’s function is thus parametrized by a single parameter *γ*_*i,i*−1_ > 0 (see below) relating the impedance mismatch between the lumped compartment and the cable [33]

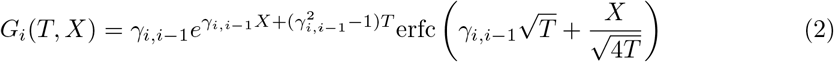

valid for *T* > 0, *X* > 0. The lumped compartment encapsulates a dependency on the state of the antidromic internode (Fig. 1 (d)). Figure 2 illustrates how this Green’s function depends on *X* and *γ*.

**Figure 2.**
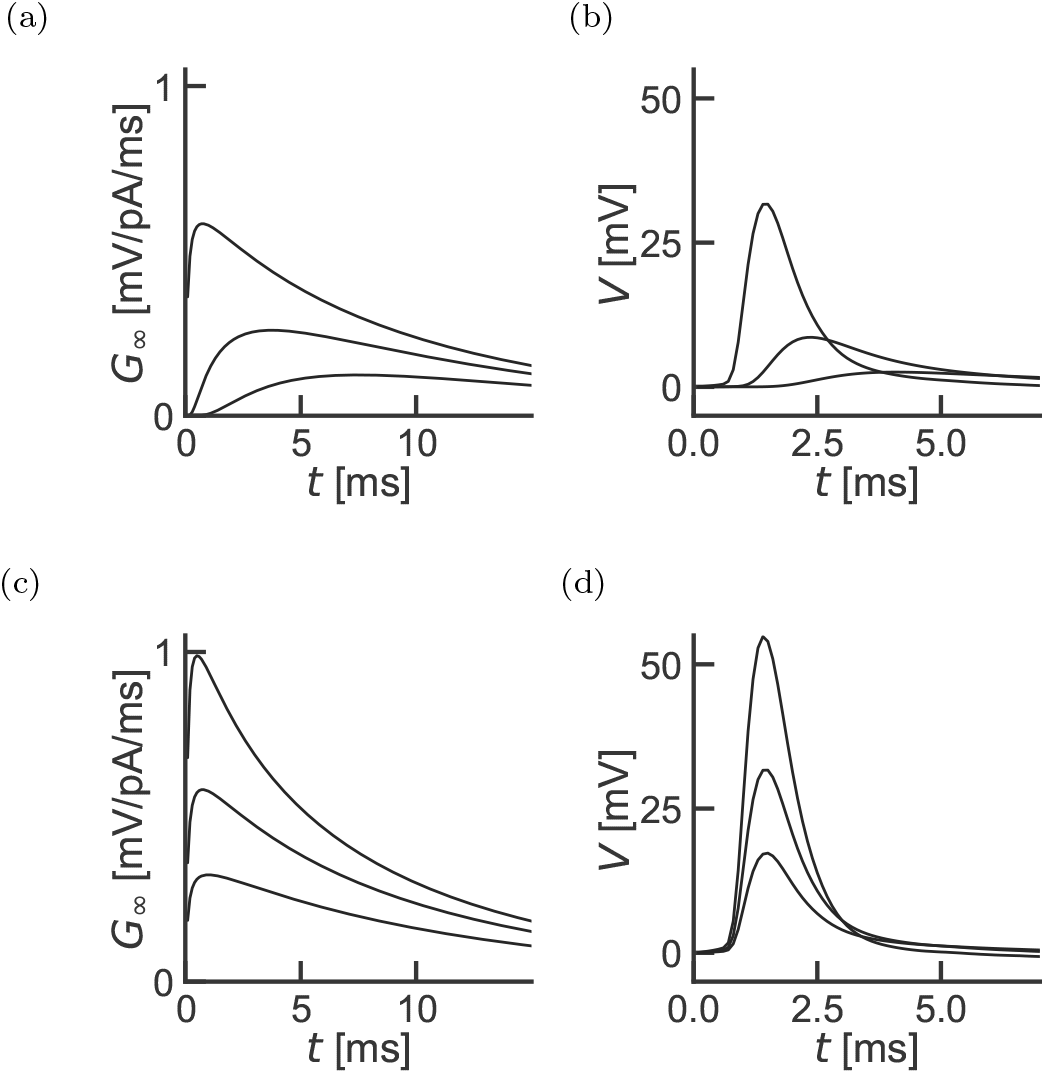
Impulse Response function for internodal current propagation. (a) The impulse-response function of a semi-infinite cable with lumped soma is shown for increasing effective distances *X* = 0.1,0.5, 1 (from top to bottom) for *γ* =1. (b) A typical action potential current filtered with the Impulse-response function shown in (a). (c) Impulse-response function of a semi-infinite cable with lumped soma for increasing electrotonic ratios *γ* = 2,1, 0.5 (from top to bottom) and *X* = 0.1. (d) Membrane potential obtained by filtering the action potential current with the Green’s functions shown in (c).

In Eq. 2, we have used *X*, the internodal distance, in units of the electrotonic length constant λ. The location of the next node is therefore the unscaled internodal distance *d* divided by the distance λ over which a steady state depolarization has fallen by a factor *e*^−1^. Since the electrotonic constant depends on the degree of myelination, we specify that the orthodromic compartment has a length constant of λ*_i_*, such that

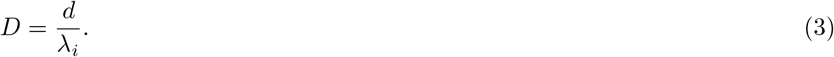

In what follows we will focus on the position of node *i* +1 using the node i as reference, *X* = *D* and the position of node *i* in the same reference system *X* = 0.

The parameter *γ* is the ratio between the total transmembrane resistance of the lumped compartment at *X* = 0 and the axial resistance *r_a_* over one electrotonic length of the orthodromic compartment *r_a_*λ*_i_* [33, 34]. If the lumped compartment has a total resistance corresponding to the sum of the axial resistance over one electrotonic length of the antidromic axon *r_a_*λ_*i*−1_ and the much smaller transmembrane resistance of the Ranvier node, *R_N_*, then we obtain the simple relation

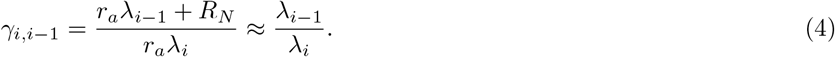

For the Green’s function of the ith internode, the parameter *γ* is the electrotonic length of the antidromic compartment λ_*i*−1_ divided by that of the orthodromic compartment λ*_i_*. For an intact axon the electronic constant of successive internodes is uniform λ_*i*−1_ = λ_i_ ≡ λ*_M_* and *γ* = 1, where λ*_M_* is the electrotonic constant of a fully myelinated internode. Similarly, for a fully demyelinated axon, the electrotonic constant is uniform and *γ* = 1. Uneven myelination will affect *γ* such that orthodromic demyelination implies *γ* > 1 (top in Fig. 1 (d)) and antidromic demyelination implies *γ* < 1 (middle in Fig. 1 (d)).

The only parameters regulating internodal filtering are, therefore, the membrane time constant *τ*, the internodal distance d and the orthodromic and antidromic electrotonic constants, λ*_i_* and λ_*i*−1_ respectively. In the following, the membrane time constant is set to a typical value *τ* =15 ms. In addition, we used an internodal distance *d* =1 mm consistent with experimental recordings [35]. The myelinated electrotonic constant is chosen to be much larger than the internodal distance to ensure rapid propagation. We used λ*_M_* = 200 mm as a baseline electrotonic constant for a fully myelinated internode.

#### Nodal Spiking

Given a membrane potential time-course from the drift-diffusion step, *V_i_*, we now calculate a probability of firing associated with this depolarizing drive. Nodal spiking is considered to be probabilistic since it results from the stochastic activation of a finite number of ion channels [36, 37, 13] amid disturbances from ephaptic coupling with neighboring axons [38, 39, 40, 41, 42]. To capture this stochasticity, we use a common approximation where all noise sources can be encapsulated in a simple mapping between a deterministic membrane potential drive *V_i_*(*T, X*) and a probability intensity, or hazard, *ρ*_*i*+1_(*T*) [43, 44, 21]. Following previous theoretical and experimental estimates of this hazard [46, 47], we assume that *ρ* is determined by an exponential function of the membrane potential. How the firing probability depends on the rate of change of the membrane potential can be included in the formalism at a later time [46]. In our model, the probability intensity is high when the membrane potential is above a fixed threshold *θ* and goes smoothly to zero when it is below. Using a scaling factor *β* we can write the probability intensity time course for the membrane potential time course *V_i_*(*T, X*) as

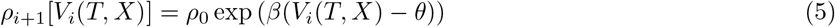

where the factor *ρ*_0_ is an arbitrary scaling constant with units of firing rate. Note that we used *V* to be the membrane potential with respect to baseline. Accordingly, the threshold *θ* is defined with respect to the same baseline. The threshold corresponds approximately to the membrane potential at which the sodium ion channels start to activate. Thus, the nonlinear relationship between membrane potential and probability of firing is thought to reflect the fact that noise may cause the neuron to fire as a function of the distance to threshold. In sum, the hazard function in Eq. 5 specifies the firing intensity of a inhomogeneous Poisson process.

This model is known as the spike-response model with escape noise [44, 21]. It has been successfully applied to experiments on cortical neurons. Given the relatively similar kinetics between action potential generation in the axon hillock and action potential generation in the Ranvier nodes, we believe that the accuracy previously shown for the soma-hillock [20] will apply also to the Ranvier Nodes.

This probability intensity is used to define the probability of transmission (Fig. 1 (c)). Using the hazard-function formalism [21], we consider the probability that a spike will have been transmitted between time zero and time *T*:

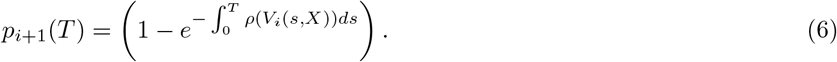

where we have omitted the dependency on *X* for simplicity of notation. We note that this expression implies a nonlinear transformation from the membrane potential time course to the probability of firing.

The probability of transmission 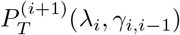 follows as might be expected from this quantity: it is the probability that a spike will have been transmitted after a sufficiently long time: 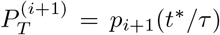. We used *t*\* = 10 ms throughout our work, which is much larger than the time taken to travel between two nodes.

To ensure a realistic dynamics, we calibrated the parameters of the model, *β, θ* and *I*_AP_, on publicly available data as follows. For the action potential time-course *I*_AP_, we used membrane potential recordings from deep cortical neurons of young rats [48] stimulated with time-dependent current-clamp input. To extract the current time course from the membrane potential time course, we computed the first derivative of the spike-triggered average. This current time course is used as a template current leaving any given internode (Fig. 1(a)). We have chosen *β* = (5mV)^−1^ to obtain a threshold variability as previously reported for human axons [13]. We chose the action potential threshold *θ* in order to obtain a physiological propagation delay given our choice of electrotonic constants, as described in the next section.

#### Internodal Delay and Speed

To determine the propagation speed, we consider a spike leaving node *i* and travelling toward the next node, *i* + 1, through internode *i*. The propagation speed will then be expressed as a function of the internodal delay and the internodal distance. Inhomogeneities in internodes *j* > *i* can in principle affect the delay involved in crossing internode i. These effects are neglected here, but they could be incorporated by using compartmental modeling to calibrate the impulse-response function for all possible states of the downstream internodes.

The membrane potential at internode *i* +1, which in our notation is the voltage at the previous node propagated over a distance *d*, i.e. *V_i_*(*T, D*(λ*_i_*)), is obtained from Eq. 1 with λ*_i_* being the electrotonic length constant of internode *i* and *γ*_*i,i*−1_ = λ_*i*−1_/λ*_i_* characterizes the impedance mismatch between internode *i* and the lumped compartment at node *i* (Eq. 4). Using the firing intensity defined in Eq. 5, we can obtain the probability that the first spike occurs at time *T* [21]

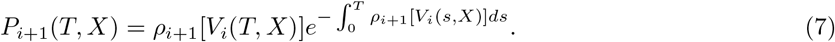

where we make explicit the dependence on *V_i_*(*T, X*) through Eq. 5.

The internodal deylay is the difference between the time at which the spike is produced at node *i*, 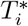 and the time at which the spike is produced at node *i* + 1, 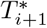, namely

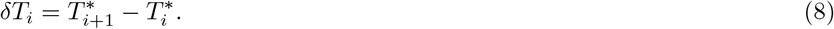

The timing of a spike at internode *i* + 1 is taken to be the maximum likelihood timing

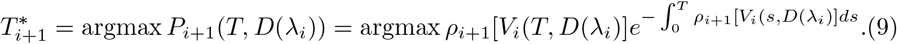

To compute the reference spike time 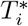, we use the membrane potential *V*_*i*−1_ calculated with *γ* = 1 in Eq. 1 and λ*_M_*:

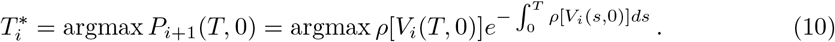

Together, these quantities are used to calculate the propagation speed v_i_ across internode *i*, that is, the internodal distance *d* divided by the internodal propagation delay

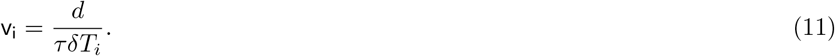

Figure 3 shows how the membrane potentials at *i* and *i* + 1 (Fig. 3(a)) lead to a propagation speed (Fig. 3(b)). This speed depends on the length constants λ*_i_* and λ_*i*−1_ as well as on the threshold *θ*. Increasing the threshold in Eq. 5 broadens the first-spike time distribution and increases the delay (Fig. 3(c-d)). For a uniform myelination with λ*_i_* = λ_*i*−1_ = λ*_M_* = 20 mm, we have *γ*_*i,i*−1_ = 1 and *X* = 0.005. The small effective distance leads to a small shift of the membrane potential at X=0 to the membrane potential at *X* = 0.005 (Fig. 3(a)). We can compute the speed for a given value of the threshold *θ*. The propagation speed v decreases as the threshold is increased (Fig. 3(b)).

**Figure 3.**
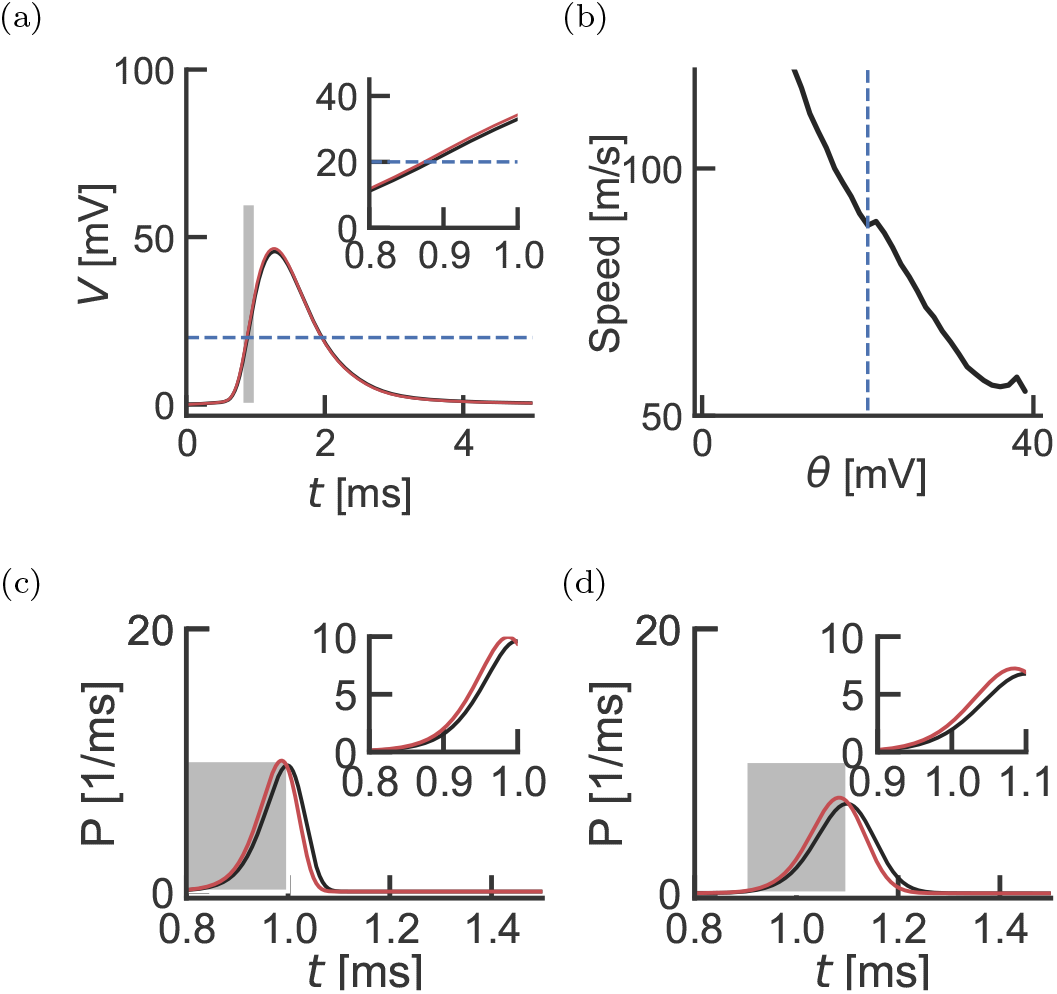
Propagation Speed. Notice that the time scales are in milliseconds. (a) Filtered action potentials at distance *X* = 0 (red) and *X* = 0.005 (black), corresponding to *d* =1 mm and λ*_M_* = 200 mm. The threshold is shown in blue. Inset expands area in the gray rectangle. (b) The resulting speed of propagation (Eq. 11) as a function of threshold *θ*. The blue dashed line indicates *θ*\*, the parameter value used in the simulations. An average transmission delay of 0.0125 ms over *d* =1 mm yields a speed of 80 m/s. (c) Difference between latency distributions calculated from the action potential at *X* = 0 and *X* = 0.005 (shown in (a)) for *θ* = 20 mV. Inset expands area in the gray rectangle. (d) Same as (c) but *θ* = 50 mV.

We use this relationship to determine a threshold that gives realistic propagation speeds. Specifically, we numerically calculate the speed v(*θ*) for every value of the threshold *θ* and find the threshold *θ*\* for which v(*θ*\*) is close to 80 m/s. This is obtained for a threshold at *θ* = 20 mV.

#### Homeostatic Threshold Compensation

In some simulations, we model a homeostatic regulation of propagation velocity by adjusting the firing threshold at every node. To do so, we numerically calculated the highest value of the threshold that would preserve the conduction velocity of v(*θ*\*) = 80 *m*/*s* within the bounds *θ* ∈ [5, 30] mV.

#### Jitter

The variability of propagation delay is quantified by the standard deviation of *δT_i_*. Accordingly, we want to estimate *σ*_*i*+1_ and *σ_i_*, the standard deviation characterizing the delay distribution *P*_*i*+1_(*T*) and *P_i_*(*T*), respectively. These standard deviations are estimated numerically from the full width at half maximum (FWHM) of *P*_*i*+1_(*T*) and *P_i_*(*T*), using *σ*_*i*+1_ = FWHM[*P*_*i*+1_]/2.35, and similarly for *σ*_0_. A Gaussian approximation on the latency distributions *P*_*i*+1_(*T*) and *P_i_*(*T*) means that the jitter associated with an interval *i* can be computed from

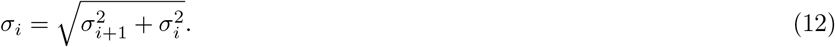

### Modeling Axonal Damage

We consider demyelinating damage. Demyelination is modeled in SSDS as an alteration to the drift-diffusion step.

#### Altered Propagation

Demyelination will affect the three principal passive cable properties differently: the relaxation time constant *τ*, the electronic length λ and the resistance ratio *γ* in Eq. 1. Firstly, the time constant is a product of transverse resistance in an internodal region *R_T_* and compartment capacitance *C, τ* = *R_T_C*. When an internode undergoes demyelination, its transverse resistance is assumed to increase while its capacitance decreases [29]. Therefore, the time constant may remain approximately constant under demyelination.

On the other hand, the electronic length is given by λ^2^ = *R_T_*/*R_a_* where *R_a_* is the axial resistance [21]. Since only the transverse resistance is affected by demyelination and not the axial resistance, the electrotonic length will decrease. Therefore, we argue that the effect of demyelination of internode is to increase the effective internodal distance through a reduction of the electrotonic length.

The resistance ratio *γ* parametrizes the effect of a mismatch between resistance of antidromic and orthodromic internodal regions. We will denote the different configurations with four cases for the value of *γ*_*i,i*−1_. *γ*_11_ corresponds to the intact axon (λ*_i_* = λ_*i*−1_ = λ*_M_*), *γ*_10_ to antidromic damage (λ_*i*−1_ = λ*_D_*), γ_01_ to orthodromic damage (λ*_i_* = λ*_D_*) and *γ*_00_ to damage on both internodes (λ*_i_* = λ_*i*−1_ = λ*_D_*). Thus, when both internodal regions are intact or when both internodal regions are equally demyelinated, we have *γ*_11_ = *γ*_00_ = 1. When only one of the internodal regions is intact, we expect *γ*_10_ < 1 for antidromic demyelination and, conversely, *γ*_01_ > 1 for orthodromic damage. Since *γ* modifies multiplicatively the current in Eq. 1, this is consistent with a greater net axial current flowing orthodromically from the Ranvier node for *γ*_01_, and conversely for *γ*_10_. Our computational work reveals in fact that the current flowing orthodromically from the spiking node will leak out of the reduced membrane resistance there, potentially making it weaker at the next node (Eq. 2). However, the current flowing antidromically will also be relatively smaller, due to the higher resistance added by the intact myelin. The result is an extra orthodromic contribution that more than compensates the first effect.

To model the effect of demyelination on the electronic length and the resistance ratio simultaneously, we suppose that the unmyelinated, or maximally demyelinated, fibre has a electronic length λ*_L_*. This will serve as a lower bound. Similarly, the intact internode has an electrotonic length λ*_M_*. A demyelinated internodal region is associated with an electrotonic length λ*_D_* between λ*_L_* and λ*_M_*. Demyelination will reduce the effective internodal distance *X* from *d*/λ*_M_* to *d*/λ*_D_*. Concurrently, it will modify the resistance ratio according to 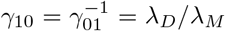, with *γ*_00_ and *γ*_11_ unchanged.

Given this parametrization of damage in terms of the electrotonic length, we quantify the intensity of damage, *D* by its relative change in *X*:

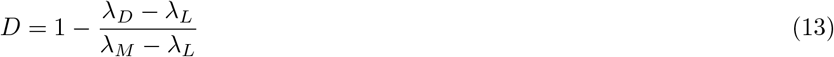

This metric, reported in percent, takes a value between 0% and 100%. We say that damage is maximal when the intensity of damage reaches *D* = 100%. We use a typical value of λ*_L_* = 1 mm for the fixed unmyelinated length constant [49]. Although modern estimates of this length constant in the cortex are about half this value [50], the results presented here do not critically depend on this lower bound. For the fully myelinated space constant we choose λ*_M_* =200 mm.

#### Delay, Failure and Jitter along a Complete Axon

We will calculate the propagation statistics in terms of four numbers: 1) the number of damaged internodes preceded by an intact internode *N*_10_, the number of damaged internodes preceded by a damaged internode *N*_00_, the number of intact internodes preceded by an intact internode *N*_11_, and the number of intact internodes preceded by a damaged internode *N*_01_. The total number of nodes is simply the sum of the number of each type of internode:

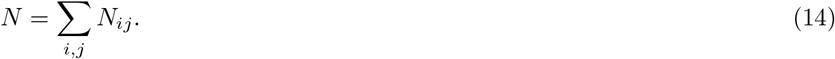

The total probability of transmission is the product of all internodal transmission probabilities:

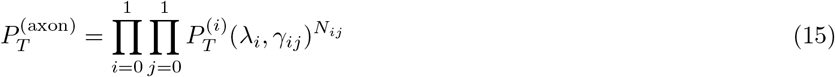

We wish to calculate the distribution of delays for a spike travelling along a complete axon, given a pattern of demyelination determined by the number of nodes in each of the four categories *N*_00_, *N*_01_, *N*_10_ and *N*_11_. Although it is possible to compute this distribution with nested convolutions of the internodal delay distributions, we will assume that each internodal delay distribution is well captured by a Gaussian. Then, the delay for the whole axon is the weighted sum of the delay associated with each type of internodes

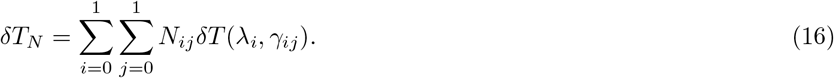

Similarly, the jitter is obtained by the weighted sum of the single internode variances.

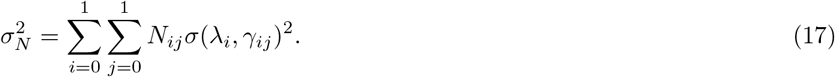

The Gaussian approximation holds well as long as the membrane potential crosses the stochastic threshold. This gives delay distributions that are sharp and approximately symmetric.

#### Modelling Damage Distribution

Here we consider a simple model where lesion can occur with constant probability *p_L_* and when a lesion occurs, it creates a damage in a fixed number, *k* =1, 2, 3,…, of successive internodes. When an internode is damaged its electrotonic length is reduced by a fixed amount that is the same for all lesions. This model is an approximation of the complex mechanisms giving rise to a correlation between the damages at subsequent internodes [51].

The parameters *k_L_* and *p* determine how the *N_ij_* are randomly generated. First we generate the number of lesions using a binomial distribution, 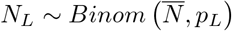:

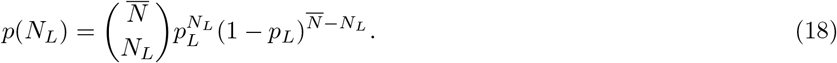

where 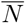 is the mean total number of internodes.

It follows that the number of nodes situated at the leading edge of a lesion *N*_01_ will equal the number of lesions, *N_L_*, unless a lesion is situated at the end of the whole axon. Similarly, the number of nodes situated at the trailing edge of a lesion *N*_10_ will equal the number of lesions *N_L_* unless a lesion is situated at the very beginning of the whole axon. For simplicity we take, *N*_01_ = *N*_10_ = *N_L_*. The number of nodes surrounded by two damaged internodes, *N*_00_, is zero if lesions consist of isolated internodes (*k* = 1). Otherwise, for *k* > 1, we have *N*_00_ = *N_L_*(*k* − 1). The number of nodes surrounded by intact internodes is then given by Eq. 14: *N*_11_ = *N* − *N_L_*(*k*+1) for *k* > 1 and *N*_11_ = *N* − 2*N_L_* for *k* =1. Here the total number of internodes *N* is not a fixed number but a function of the random number *N_L_* (Eq. 14). In the regime *k* << *N* and *p_L_* << 1, the fluctuations of *N* around 〈*N*〉 will be relatively small. The average number of lesion that results from this scenario is 〈*N_L_*〉 = 〈*N*〉_*p_L_*_.

The overall *temps perdu*, Δ*TP*, averaged over the lesion configuration thus becomes a function of the average number of 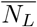,

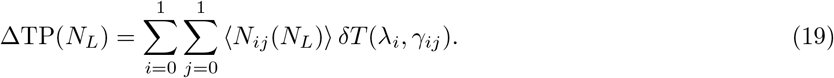

and similarly for the variance,

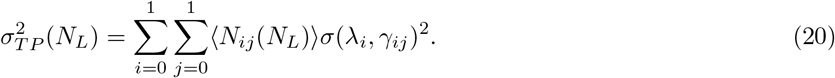

Together Eqs. 19–20 describe a Gaussian approximation to the spike time distribution at the end of an axon.

### Compound Action Potential

Previous theoretical investigations [52, 53] have concluded that the local field potential (LFP) can be well captured by monopole terms for the current sources. For myelinated nerves, current sources consist mostly of the transmembrane currents underlying the action potential generation at the nodes of Ranvier. The LFP is then obtained by summing the contributions of each node of Ranvier in the proximity to the recording electrode. Given a relatively large internodal distance, only the nodes closest to the recording electrode will contribute to the LFP. We can therefore sum for each axon l only the contribution of the nearest node. Let the electrode be closest to the nth node of each axon, and write the current of the nearest node as *I_ln_*(*t*) and its location with respect to the recording electrode as *r_ln_*. We have the electric potential *ϕ* arising from transnodal currents scaled by a Coulombic factor:

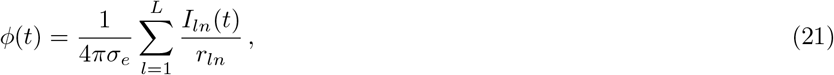

where *σ_e_* is the electric permittivity [52]. Equation 21 can be related to our framework by considering that the current at time t in a given node is the action potential current triggered with a delay *δT_In_*: *I_ln_*(*t*) = *I_AP_* (*t* − *δT_ln_*).

According to the law of large numbers, the sum of the distance-scaled nodal currents over a large number of independent and identically distributed currents should be close to their expected value. Also, since we are only interested in the relative amplitude of the compound action potential, we lump the scaling factors into a constant c and focus on the time-dependence. This scaling constant captures geometrical factors which will scale the LFP uniformly depending on the spatial distribution of the nearest nodes. We therefore obtain

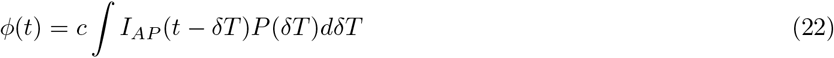

where we have dropped the subscript n for simplicity. The compound AP is proportional to the convolution between the cross-membrane current during an action potential and the latency distribution of the nearest internode.

We apply the central-limit theorem to obtain the following form for the compound action potential:

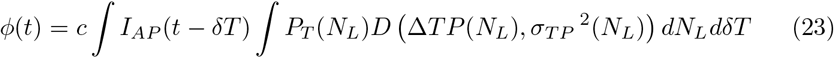

where *D*(*μ, σ*^2^) is a normal distribution with mean *μ* and variance *σ*^2^. The mean and the variance are given by Eq. 19 and Eq. 20, respectively.

Following the Gaussian approximation of Eq. 16, the distribution of delays for propagation between node 0 and node N can be written as

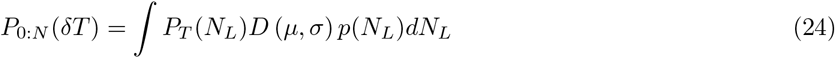

and the number of lesions follows a binomial distribution (Eq. 18). Taken together, Eqs. 23–18 determine the compound action potential. The main quantities of interest are listed in Table 1 below. A summary of the steps used to compute the probability of propagation, jitter distributions and compound action potential is given in the first part of the Results section.

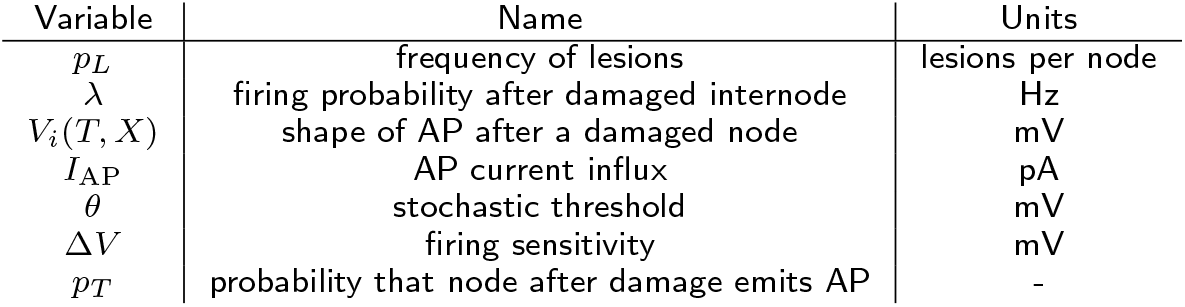

## Results

In order to study how demyelination affects action potential transmission over long axons, we constructed a computational model that captures the essential features of saltatory propagation. The stochastic spike-diffuse-spike model (SSDS: see Methods) iterates between a stochastic firing step and a cable-filtering step. In the drift-diffusion step, firing in node *i* causes a membrane potential change in node *i* + 1 as determined by the cable impulse response function. This step modifies the width and the amplitude of the action potential in a manner phenomenologically equivalent to drift-diffusion. In the firing step, we determine the firing probability given the membrane potential reaching this node. The biophysical mechanisms underlying stochastic firing are not specified in the SSDS but they are thought to reflect both the stochastic opening of possibly distinct types of voltage-dependent ion channels in the node of Ranvier [36, 37, 45, 13] and ephaptic couplings with neighboring axons [38, 39, 40, 41, 42]. This SSDS model allows us to estimate transmission probability, propagation delay and spike timing jitter for any axon length as a function of the number of damaged nodes.

Figure 1 illustrates the SSDS model for three distinct local patterns of demyelination for a spike leaving a given node and propagating through the orthodromic internode to the next node. Upon activation of the first node of Ranvier, the stereotypical time-course of the action potential is observed (Fig. 1 (a)). Depending on the demyelination pattern, the stereotypical current from the first node will produce a depolarization in the next node (Fig. 1 (b)). This depolarization time course is used to calculate the probability that an action potential is produced in this node (Fig. 1 (c)). Three distinct types of demyelination pattern are distinguished (Fig. 1 (d)): demyelination of the orthodromic internode only, demyelination of the antidromic internode only, and demyelination of both antidromic and orthodromic regions. If a spike is generated in the next node, the process continues along the axons iteratively.

Parameters of the SSDS consist of a non-parametric time-dependent current resulting from the stereotypical time-course of ionic currents in the Nodes of Ranvier (*I*_AP_), the impulse-response function regulating propagation of this current to the next node of Ranvier (*G_X_*), the firing threshold at the nodes (*θ*) and the stochastic firing sensitivity (*β*). Parameter values are either calibrated using published experimental data or follow classical modeling studies (see Methods). Briefly, first we estimate *I*_AP_ from the spike-triggered average membrane potential from patch-clamp recordings. We note that although the framework is general and can take into account the spike after-hyperpolarization [4], we have restricted the present analysis to time scales shorter than 10 ms. Second, the impulse response function is parametrized according to the theoretical treatment of Rall [33] where a lumped compartment corresponding to the node and its antidromic internode is followed by a semi-infinite uniform cable (Fig. 2) corresponding to the orthodromic internode. Then, both the firing threshold and sensitivity are chosen in order to obtain standard values of propagation speed and threshold variability (Fig. 3).

Demyelination will directly affect directly the impulse-response function. In fact we find that for our parametrization, two parameters regulate the shape of the impulse-response function. For a node *i* followed by internode i and preceded by internode *j*, these two parameters are the electrotonic length constant for the orthodromic cable λ*_i_* and the ratio of orthodromic and antidromic length constants *γ_ij_*. Demyelination reduces the length constant by an amount reflecting the degree of demyelination (see Methods). Figure 2 illustrates how the different filtering properties arise from changes in the electrotonic constant λi keeping the antidromic one fixed. Both the impulse-response function amplitude and temporal profile are affected by changes in the orthodromic length constant (Fig. 2 (a)). As a result, a decrease in λ*_i_* will produce both a dampening and a broadening of the depolarization reaching the next internode (Fig. 2 (b)). In contrast, modifying the antidromic-to-orthodromic ratio of length constants *γ_ij_* while keeping the orthodromic electrotonic length fixed modifies the amplitude of the impulse-response function without affecting its time-course (Fig. 2 (c)). This results in a dampening of the depolarization reaching the next node, which is seen without an associated broadening (Fig. 2 (d)). Therefore our SSDS model is biophysically justified, and it can be used to understand how different demyelination patterns relate to propagation properties.

### Demyelination and Propagation Properties – Single Internode

To study the effects of demyelination on single internodal propagation, we derive expressions for propagation delay (Eq. 19), spike timing jitter (Eq. 20) and transmission probability (Eq. 15) in the SSDS model. We then study the effect of demyelination intensity for the three different damage patterns illustrated in Fig. 1 (d), namely damage affecting the orthodromic region, the antidromic region, or both regions with equal intensity. The degree of demyelination is quantified with the metric *D* (see Methods, Eq. 13), which varies between 0% for intact internodes, and 100% for complete demyelination.

We find that transmission probability is only affected at extreme damage intensities for the uniform and orthodromic damage patterns (Fig. 4 (a)). This is consistent with previous computational studies showing that a very high degree of demyelination is required to prevent internodal propagation [2, 54, 55]. When the damage is antidromic, however, the probability of transmission starts to drop sharply at mild damage intensities (*D* = 60%). In the SSDS model, this effect results from the impedance imbalance, where an antidromic demyelination reduces the impedance in the antidromic direction. This reduces the net axial current flowing in the orthodromic direction and therefore prevents propagation. Consistent with this view, the delay and jitter both increase for weak demyelination intensities when the damage is antidromic (Fig. 4 (b)-(c)). In the case of uniform damage, increases in both the delay and jitter are observed only for very large demyelination intensities.

**Figure 4.**
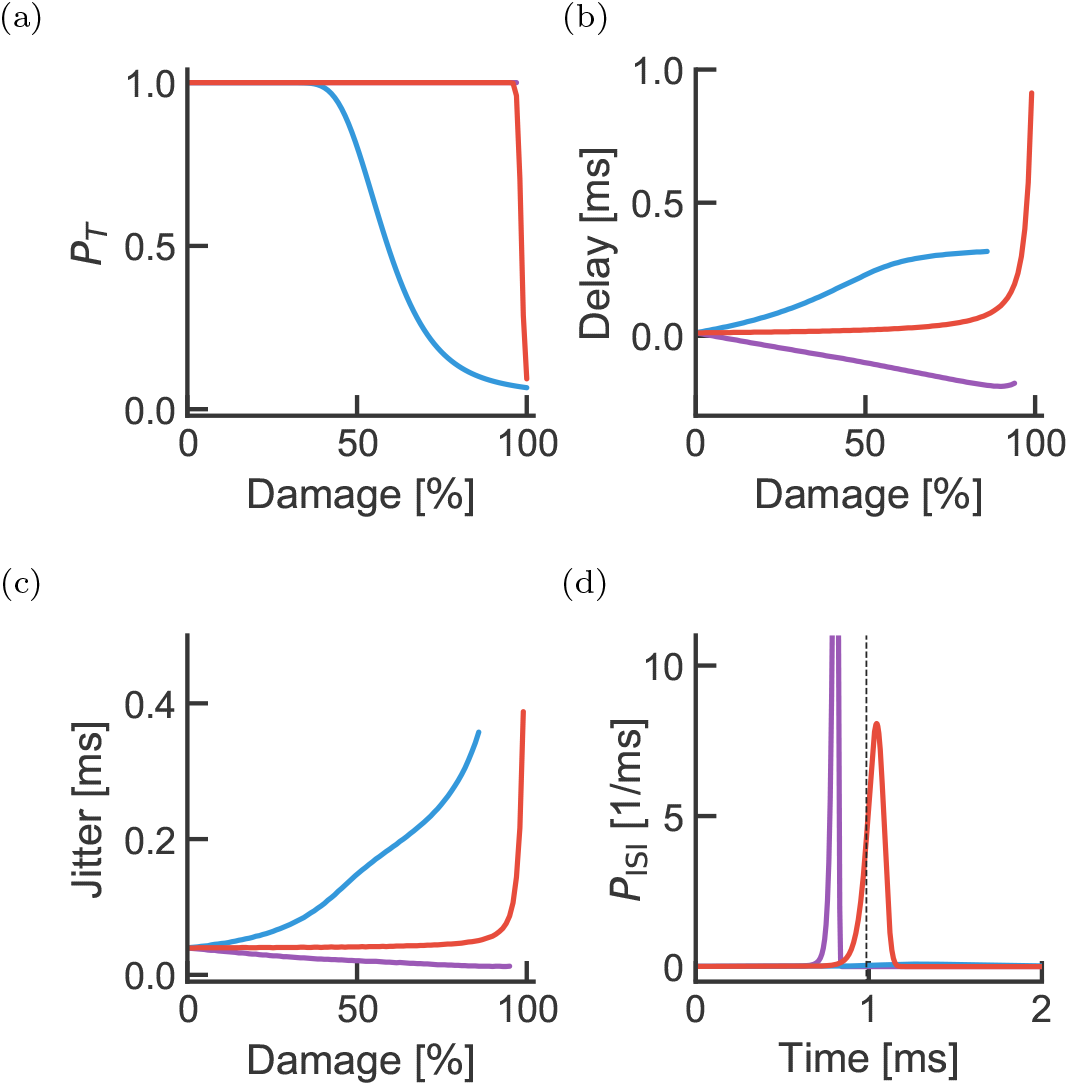
Effect of demyelination on propagation across a single internode. (a) Effect of demyelination on the probability of transmission *p_T_* for the three cases shown in Fig. 1 (purple is orthodromic damage, blue is antidromic damage and red is damage in on both side of the node). Purple line is hidden behind the red curve, but stays at *p_T_* = 1 over the range of damage intensities studied. (b) Effect of demyelination on transmission delay. Negative delays means a propagation delay shorter than in the absence of damage. Delays corresponding to *p_T_* < 0.1 are not plotted. (c) Same as (b) but for transmission jitter. (d) Spike timing distributions corresponding to 80% damage (based on a time discretization of 0.01 ms).

In contrast, when the damage is orthodromic, the impedance mismatch forces more current to flow to the orthodromic side, which compensates for the reduced length constant. This leads to an acceleration of the propagation (Fig. 4 (b)) and a decrease in the jitter (Fig. 4 (c)). These results suggest unexpectedly that propagation is more likely to fail when an action potential attempts to leave a demyelinated region following a healthy region than when it attempts to traverse a demyelinated region after successfully entering it. Also, at the weak demyelination intensities where propagation is preserved (*p_T_* > 0.8, *D* < 50%), mild damage should cause a net increase in spike timing jitter whereas the net delay does not vary strongly with damage.

### Lesion Patterns and Propagation Properties – Whole Axon

We now turn to the demonstration of the power of the model to predict whole axon propagation properties. To determine how demyelination patterns affects propagation along the length of an axon, we consider demyelination in lesions made of segments of *k* contiguous internodes. Lesions are then assumed to arise randomly with probability *p_L_*. Transmission probability, propagation delay and spiking jitter are then computed analytically (see Methods). For a fixed, mild degree of demyelination, increasing the lesion probability sharply decreased the probability of transmission over long axonal distances (*D* = 50%, *k* = 1, Fig. 5 (a)). The total transmission delay increases linearly with distance, consistent with a constant but damage-dependent speed of propagation. Lesion probability strongly affects this effective propagation speed (Fig. 5 (b)). In contrast, the jitter increases sub-linearly with the number of internodes travelled (Fig. 5 (c)), consistent with a linear increase in variance. These results suggest that lesion probability strongly affects propagation properties even for a mild degree of demyelination.

**Figure 5.**
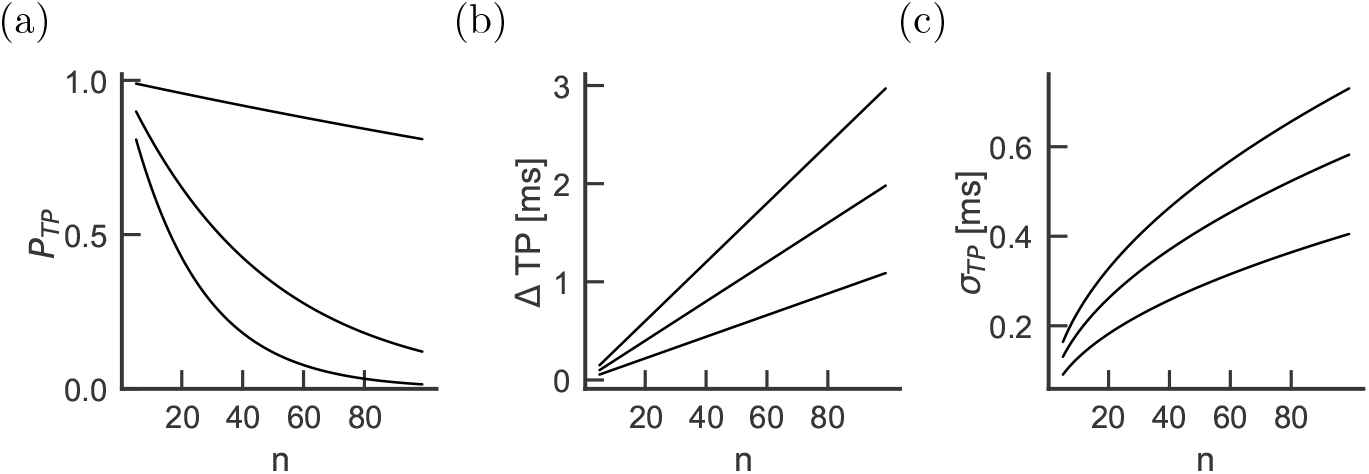
Effect of lesion frequency as a function of the number of successive internodes. For a fixed lesion size *k* = 1 and damage intensity *D* = 50%, the probability of transmission (a), the transmission delay (b) and the jitter (c) are shown as a function of the average total number of internodes. Lesion frequencies are plotted for *p_L_*=0.01, 0.1, 0.2 (from bottom to top in panel a and from top to bottom in panels b and c).

The effects of lesion probability can be contrasted with the effects of lesion size. In Figure 6 we vary the lesion size *k* and plot the distance dependence of transmission probability, propagation delay and spike time jitter. Interestingly, transmission probability is mostly unaffected by lesion size. This can be explained by noting that, in our model, increasing lesion size increases the proportion of nodes with damage both in the orthodromic and antidromic internodes (Fig. 1, red) without affecting the proportion of asymmetric damage configurations. Since symmetric damage configuration affects propagation properties only at very strong demyelination degree (Fig. 4, red curves), lesion size does not affect propagation properties at weak demyelination intensities.

**Figure 6.**
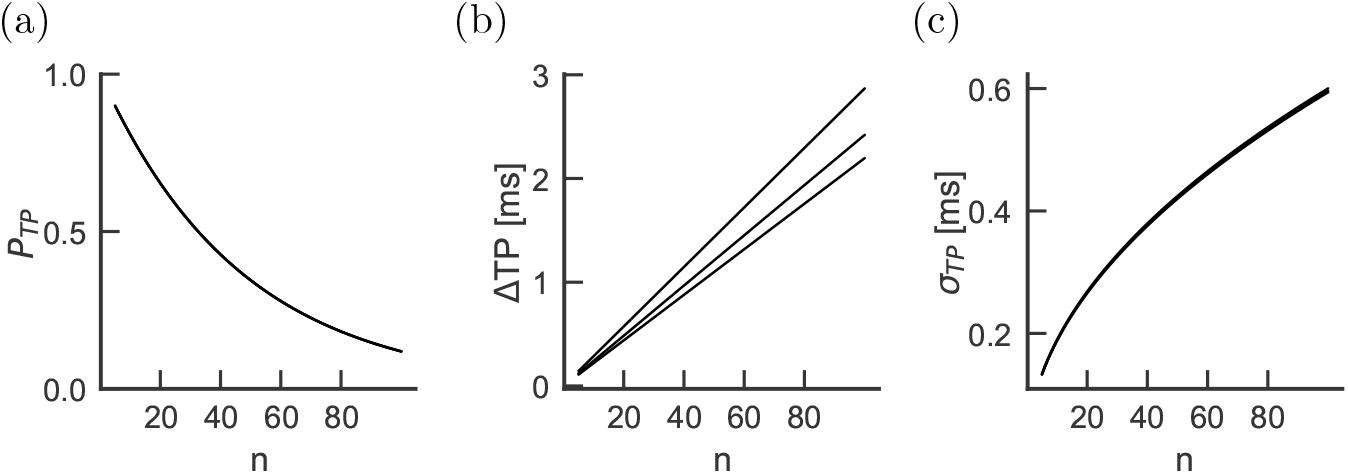
Effect of lesion size as a function of the number of successive internodes. The setup is the same as for Figure 5, but for three lesion sizes: *k* = 1, 3, 7 (from bottom to top) and fixed lesion frequency *p_L_* = 0.1. Note that the probability of transmission and the jitter depend very weakly on lesion size, as the lines perfectly overlap.

### Homeostatic Control of Demyelination-Induced Delays

Given the substantial loss of propagation produced by mild demyelination intensities in the antidromic case (Fig. 4 (a)), it appears surprising that propagation appears to be preserved amid evident disorder associated with demyelinating diseases. Nodal remodeling, particularly of axonal excitability, is believed to play an important role in preserving axonal conduction [56, 57] and may underlie the periods of remission in multiple sclerosis or other demyelinating diseases [58]. An increase of the density of sodium ion channels was linked to an improved propagation in demyelinated axons [2] and to membrane reconfiguration following demyelination [59, 60]. Increasing the density of sodium ion channels produces a lower action potential threshold [61]. This suggests a homeostatic control mechanism where the reduced transmission due to weaker depolarization (illustrated in Fig. 1 (b)) is compensated by a lower threshold. Alternatively, when demyelination enhances the depolarization, the node would ideally raise its spiking threshold so as to preserve the propagation speed. In order to model this homeostatic adjustment, we have replaced the fixed firing threshold *θ* by a threshold adjusted to be the highest value that would preserve conduction velocity. This homeostatic compensation is limited in the model by restricting the adjustment range for the threshold (see Methods, Section).

We start by considering the effect of homeostatic compensation on the propagation properties of a single internode. As expected from the observation that delays increase (decrease) when damage is antidromic (orthodromic) (Fig. 4), the compensated threshold was low for the antidromic damage configuration, unchanged for the symmetric damage configuration and increased for the orthodromic damage configuration (Fig. 7 (a)). This homeostatic compensation ensured a high transmission probability even for severe demyelination. Delays and jitters depend on damage configuration and intensity, in a manner that remains qualitatively identical to the absence of threshold compensation (compare Fig. 7 (c)-(d) with Fig. 4 (b)-(c)).

**Figure 7.**
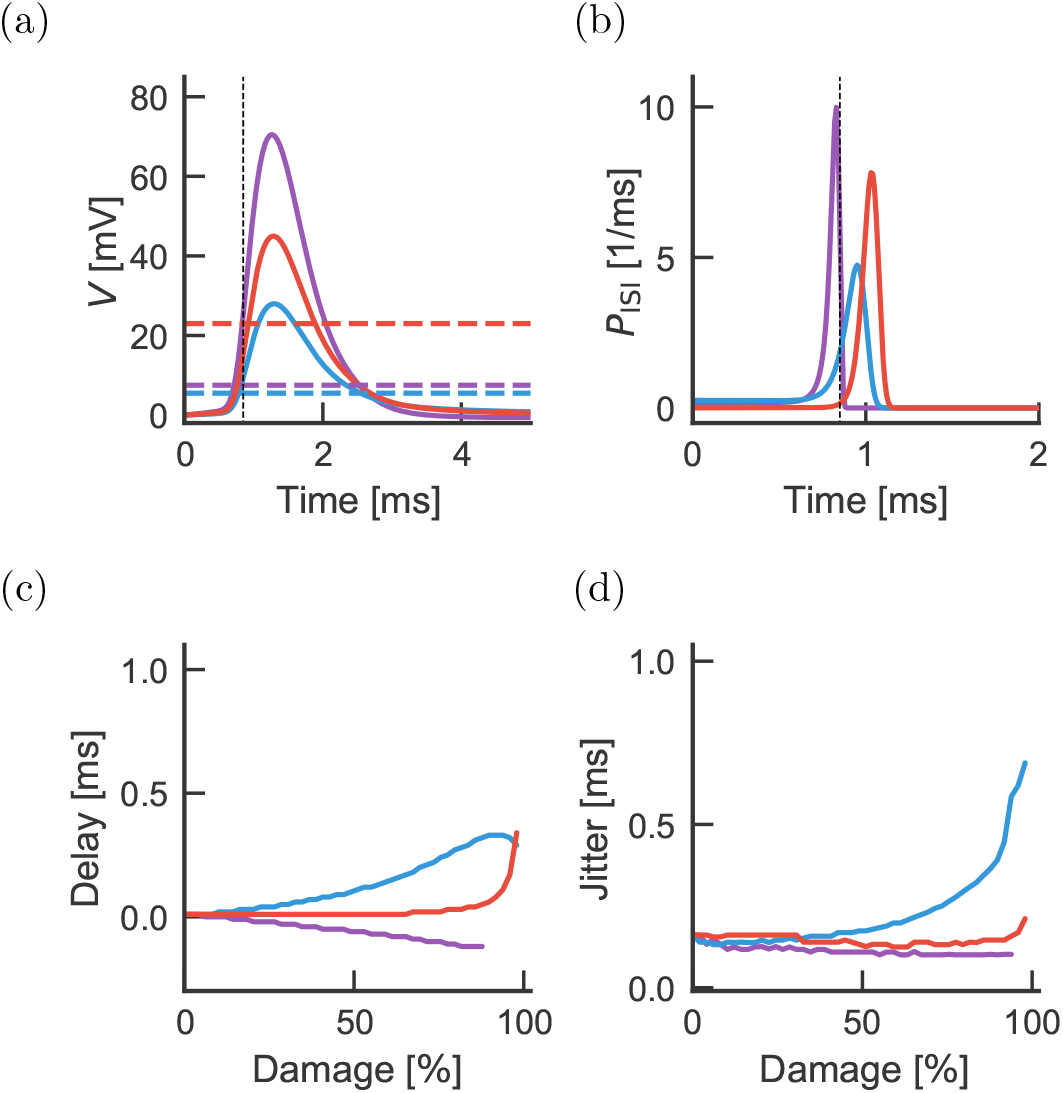
Homeostatic adjustment of threshold partially compensates for demyelination-induced delays and transmission probability across a single internode. For each demyelination intensity and depending on the antidromic-orthodromic location of the damage, we attempted to recover undamaged delays by adjusting the spiking threshold. (a) For *D* = 40%, the membrane potentials reaching the next node of Ranvier (full lines) are shown with their corresponding firing thresholds that minimize delay changes (dashed horizontal lines). (b) Delay probability distributions for the membrane potentials shown in (a). (c) The residual delay and (d) spike timing jitter are shown as a function of damage intensity. Colours follow the convention in Fig. 1.

However, quantitatively the picture is very different. As imposed by homeostatic compensation, the delay across a single internode remained approximately constant for a larger range of damage intensities (Fig. 7 (c)-(d)). As the degree of demyelination increases, the change in threshold fails to compensate for this damage, with the result that both delay and jitter are affected. Importantly, we note that, at mild antidromic damage (*D* = 50%), although the delay falls from 0.2 ms (Fig. 4 (b)) to 0.1 ms (Fig. 7 (c)) when introducing threshold compensation, the jitter remains close to 0.2 ms without (Fig. 4 (c)) and with (Fig. 7 (d)) threshold compensation. These observations suggests that the measurement of jitter could reveal demyelination, whether homeostatic compensation were at work or not.

To determine the effects of compensated demyelination on the whole axon, we determine transmission probability, delay and jitter as a function of the total number of nodes (Fig. 8). Similar to the case without homeostatic compensation, the propagation properties depend only weakly on the lesion size. We therefore focus on the effects of lesion probability. For mild damage intensities (*D* = 50%), transmission is preserved even when lesion probability is 0.2 (Fig. 8 (a)). Propagation speed decreases with lesion probability, but delays remain 2-3 fold smaller than in the absence of threshold compensation (Fig. 8 (b)). In comparison, although larger without compensation (Fig. 4 (c)), remain of the same order of magnitude as with compensation (Fig. 7 (d)). *We conclude that spike timing jitter can be a good predictor demyelination frequency and intensity even when there is nodal excitability compensation*.

**Figure 8.**
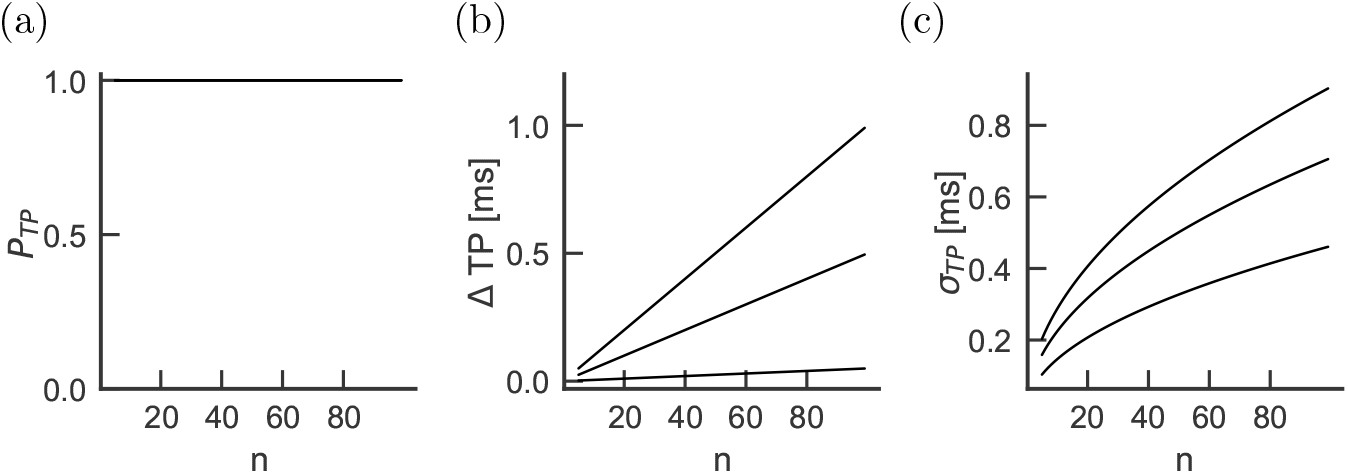
Effect of lesion frequency over multiple internodes in the presence of homeostatic compensation. The probability of transmission (a), the transmission delay (b) and the jitter (c) are shown as a function of the number of internodes through which the activity propagates. Three lesion frequencies are plotted: *p_L_* =0.01, 0.1, 0.2 (from bottom to top in panels b and c, perfect overlap in panel a). The demyelination of each lesion corresponds to 50% damage.

### Compensated Demyelination can be inferred from the Compound Action Potential

To investigate whether the effects of demyelination may be observed experimentally, we estimated the extracellularly recorded compound action potential expected from a bundle of simultaneously activated axons (see Methods, Eq. 23). We illustrate our results in the context of homeostasis. We study the hypothesis that an increase in jitter, by reducing the synchrony within the bundle, may affect the width of the compound action potential. Figure 9 (a) shows that mild (*D* = 50%) lesion probability increases both the delay of the first trough and the width of the compound action potential. The propagation is strongly impeded by rare but severe (*D* = 97%) lesions as revealed by the amplitude of the compound action potential (Fig. 9 (b)). Figure 9 (c) shows how the full width at half maximum (FWHM) has a clear dependence on both the damage intensity D and the lesion frequency *p_L_*. We conclude that the width of the compound action potential, like the jitter of an action potential along a single axon (Fig. 8 (c)), can be a good predictor of demyelination frequency and intensity even in the presence of nodal excitability compensation.

**Figure 9.**
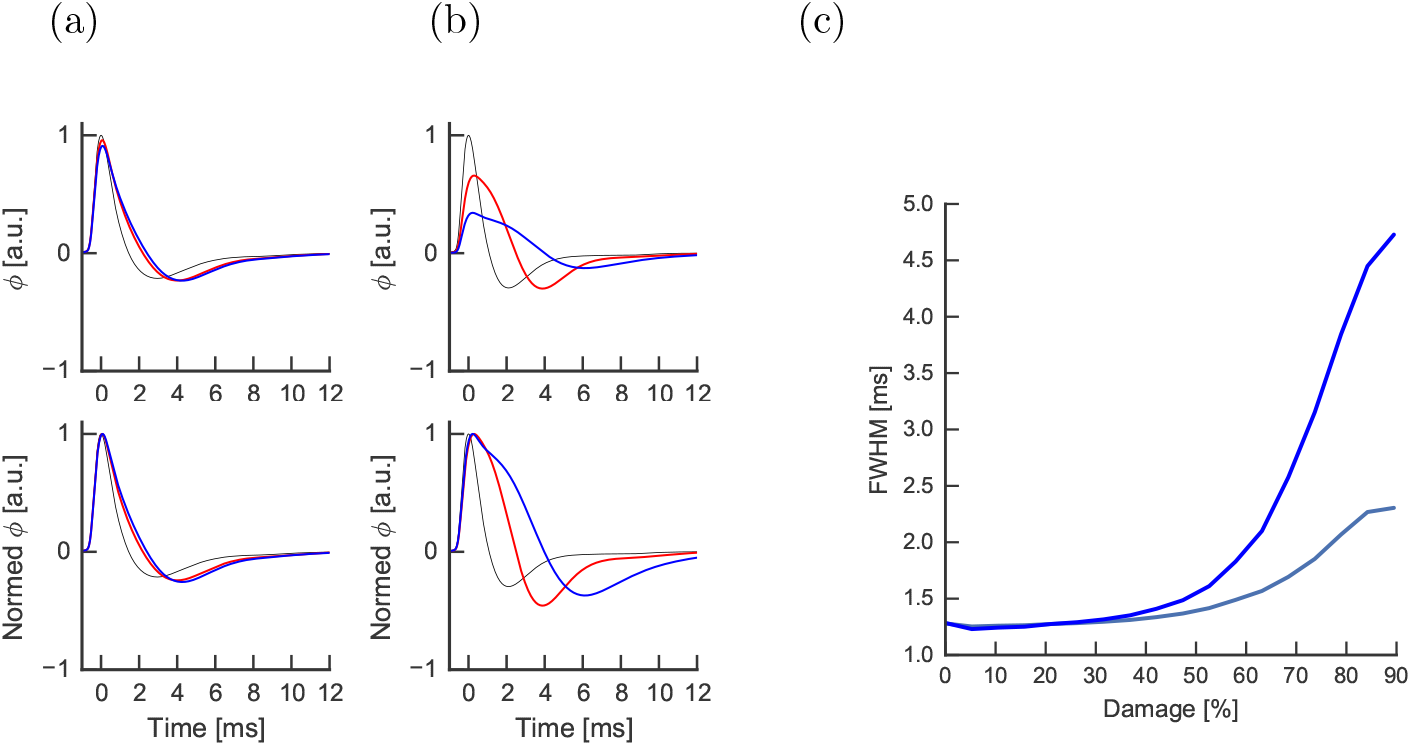
Effect of lesion size and intensity on the compound action potential in the presence of threshold compensation. The compound action potential is shown (top) for an intact axon with 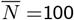 internodes (black line) and two lesion frequencies *p_L_* = 0.01 (red) and *p_L_* = 0.2 (blue). Intensity of damage and lesion size are kept fixed (*D* = 50%, *k* = 1). The time course is normalized to the peak amplitude of intact axons (bottom). (b) Compound action potential for changing lesion intensity *D* = 70% (red) and *D* = 97% (blue) while keeping size and frequency of lesions fixed (*p_L_* = 0.1, *k* = 1). (c) The Full Width at Half Maximum (FWHM) of the compound action potential as a function of damage intensity for rare and small lesions (gray, *p_L_* = 0.1, *k* =1) and frequent but small lesions (blue, *p_L_* = 0.2, *k* = 1).

## Discussion

We have presented a modeling framework to estimate propagation delay and jitter in weakly or sporadically demyelinated axons. The framework extends the SSDS model of propagation along an unmyelinated dendrite with equally spaced excitable spines [25, 62, 63] to the context of a myelinated axon with excitable nodes of Ranvier. Our goal is to account for different spike shapes and stochastic firing effects. The simplified computational model can be used to simulate propagation over a very large number of nodes and internodes, and also to determine how weak or sporadic damage can cumulate over large distances in terms of probability of transmission and the mean and variance of the propagation speed and delay. Waxman (1976) studied the effect of damage distribution, but without focusing of the cumulative effect of transmission probabilities, delays and jitter. Our use of a reduced description allows us to investigate axons of realistic, and in fact arbitrary length [65].

Our main findings can be summarized as follows. First, we found that orthodromic damage must be strong to affect the transmission probability. Also, antidromic damage has a more significant effect when a spike leaves a demyelinated region and goes into an myelinated region. As a result, propagation delay down a sequence of internodes can be increased or decreased. Since a spike traversing a demyelinated region first faces damage in an orthodromic compartment and then in an antidromic compartment, the decreased delay is followed by a increase delay, with a net increase of the delay. If homeostatic compensation reduces how damage changes the propagation delay, we see that damage is revealed by propagation jitter and almost invisible from the propagation delay. Finally, we show that this effect can be observed from the dispersion of the compound action potential.

We will discuss two particular findings, namely the effect of the antidromic impedance mismatch and the effect of demyelination on the compound action potential and spike timing jitter.

Early computational studies have shown that the action potential has a lower amplitude in both the node before and the node after a partially demyelinated internode [9]. Changes in impedance can result in conduction block even when the demyelinated region contains sodium and potassium ion channels [2]. Our simulations indicate that an additional delay is produced when the action potential leaves a partially demyelinated internode and enters into an intact region. These results are consistent with compartmental simulations of mild paranodal damage [5], but further work will be required to test experimentally our predictions regarding jitter and delay upon leaving demyelinated regions. In particular, this result relies on the fact that in our model, nodal current between two intact internode will flow in equal proportion in orthodromic and antidromic compartments because the current is divided equally between compartment with matched impedance. This situation may be affected by the mismatch of orthodromic and antidromic potential during propagation. Detailed simulations of a demyelinated segment can resolve this issue.

Our estimates of the compound action potential show that its FWHM can triple for rare but intense demyelination. This is consistent with previous experimental recordings of pharmacological demyelination [26], and in patients with demyelinating diseases [27, 28]. Our result that either an increase or a decrease in propagation delay can result from a damaged internode could explain why the compound action potential delay itself may be a less potent parker of demyelination [66, 67, 35] than the compound action potential dispersion.

The approach we have used relies on five critical assumptions delineated in Methods, of which we discuss three here. First, we have assumed a specific form of the linear impulse-response function based on a classic theoretical derivation for a uniform semi-infinite cable with lumped compartment. Many properties of the axon may affect this function. Some of these characteristics may be reasonably captured by the changes in electrotonic length constants considered here, but in other cases an different modulation of the impulse response function must be considered. Ideally, experimental measurements would either shore up or replace the hypotheses used here.

Second, we have not considered the active propagation along demyelinated internodes mediated by an increase in sodium and potassium ion channels in the demyelinated region [35]. Although a subthreshold activation of these active conductances can be treated in the spike-diffuse-spike framework [62], it cannot capture a fully nonlinear dynamical response of the internodes. These two assumptions imply that the modelling framework should be reserved for the study of relatively mild demyelination where ion channels remain in the nodes of Ranvier and saltatory propagation is preserved.

Finally, implicit to the SSDS model, we assumed that there can be no interactions between nodes that are more than two internodal regions apart. Triplet or quadruplet interactions can be taken into account, but would require additional extensions of the modeling framework. Also, the same formalism with the same caveats could be applied to other situations such as the study of traumatized axons [68]. As a promising next step, our theoretical framework could be used in combination with combined measures of clinical scores and demyelination properties [69] to identify the main impediments to propagation as well as the most effective markers of weak demyelination.

## Author’s contributions

AL and RN conceived the study and wrote the article. RN developed the theoretical framework and analyzed the results.

## Acknowledgements

RN was supported by an NSERC Discovery Grant (RGPIN-2017-06972) and the Canadian Neurophotonics platform. AL was support by NSERC Discovery Grant (RGPIN-2014-06204). We thank Daniel C. Cote and Yves De Koninck for helpful discussions.

## References

1. Helmholtz, H.: Note sur la vitesse de propagation de l?agent nerveux dans les nerfs rachidiens. CR Acad Sci (Paris) 30, 204–206 (1850)

2. Waxman, S.G., Brill, M.H.: Conduction through demyelinated plaques in multiple sclerosis: computer simulations of facilitation by short internodes. Journal of Neurology, Neurosurgery & Psychiatry 41(5), 408–416 (1978)

3. Waxman, S.G., Wood, S.L.: Impulse conduction in inhomogeneous axons: effects of variation in voltage-sensitive ionic conductances on invasion of demyelinated axon segments and preterminal fibers. Brain research 294(1), 111–122 (1984)

4. McIntyre, C.C., Richardson, A.G., Grill, W.M.: Modeling the excitability of mammalian nerve fibers: influence of afterpotentials on the recovery cycle. Journal of neurophysiology 87(2), 995–1006 (2002)

5. Babbs, C.F., Shi, R.: Subtle paranodal injury slows impulse conduction in a mathematical model of myelinated axons. PLoS One 8(7), 67767 (2013)

6. FitzHugh, R.: Impulses and physiological states in models of nerve membrane. Biophys. J. 1, 445–466 (1961)

7. Basser, P.: Cable equation for a myelinated axon derived from its microstructure. Medical & biological engineering & computing 31(1), 87–92 (1993)

8. Nygren, A., Halter, J.: A general approach to modeling conduction and concentration dynamics in excitable cells of concentric cylindrical geometry. Journal of theoretical biology 199(3), 329–358 (1999)

9. Koles, Z., Rasminsky, M.: A computer simulation of conduction in demyelinated nerve fibres. The Journal of physiology 227(2), 351 (1972)

10. Blight, A.: Computer simulation of action potentials and afterpotentials in mammalian myelinated axons: the case for a lower resistance myelin sheath. Neuroscience 15(1), 13–31 (1985)

11. Halter, J.A., Clark Jr, J.W.: A distributed-parameter model of the myelinated nerve fiber. Journal of theoretical biology 148(3), 345–382 (1991)

12. Stephanova, D.I., Daskalova, M.S., Alexandrov, A.S.: Differences in membrane properties in simulated cases of demyelinating neuropathies: Internodal focal demyelinations with conduction block. Journal of biological physics 32(2), 129–144 (2006)

13. Hales, J.P., Lin, C.S.-Y., Bostock, H.: Variations in excitability of single human motor axons, related to stochastic properties of nodal sodium channels. J Physiol 559(3), 953–964 (2004)

14. Zeng, S., Jung, P.: Mechanism for neuronal spike generation by small and large ion channel clusters. Physical Review E 70(1), 011903 (2004)

15. Zeng, S., Tang, Y., Jung, P.: Spiking synchronization of ion channel clusters on an axon. Physical Review E 76(1), 011905 (2007)

16. Ochab-Marcinek, A., Schmid, G., Goychuk, I., Hänggi, P.: Noise-assisted spike propagation in myelinated neurons. Physical Review E 79(1), 011904 (2009)

17. Pillow, J., Paninski, L., Uzzell, V., Simoncelli, E., Chichilnisky, E.: Prediction and decoding of retinal ganglion cell responses with a probabilistic spiking model. Journal of Neuroscience 25(47), 11003–11013 (2005)

18. Pillow, J., Shlens, J., Paninski, L., Sher, A., Litke, A., Chichilnisky, E., Simoncelli, E.: Spatio-temporal correlations and visual signalling in a complete neuronal population. Nature 454(7207), 995–999 (2008)

19. Mensi, S., Naud, R., Avermann, M., Petersen, C.C.H., Gerstner, W.: Parameter extraction and classification of three neuron types reveals two different adaptation mechanisms. Journal of Neurophysiology 107, 1756–1775 (2012)

20. Pozzorini, C., Naud, R., Mensi, S., Gerstner, W.: Temporal whitening by power-law adaptation in neocortical neurons. Nat. Neurosci. 16(7), 942–948 (2013)

21. Gerstner, W., Kistler, W., Naud, R., Paninski, L.: Neuronal Dynamics. Cambridge University Press, Cambridge (2014)

22. Naud, R., Bathellier, B., Gerstner, W.: Spike-timing prediction in cortical neurons with active dendrites. Front. Comput. Neurosci. 8, 90 (2014)

23. Pozzorini, C., Mensi, S., Hagens, O., Naud, R., Koch, C., Gerstner, W.: Automated high-throughput characterization of single neurons by means of simplified spiking models. PLoS Comp. Biol. 11(6), 1004275 (2015)

24. Teeter, C., Iyer, R., Menon, V., Gouwens, N., Feng, D., Berg, J., Szafer, A., Cain, N., Zeng, H., Hawrylycz, M., et al.?. Generalized leaky integrate-and-fire models classify multiple neuron types. Nature communications 9(1), 709 (2018)

25. Bressloff, P.C., Coombes, S.: Synchrony in an array of integrate-and-fire neurons with dendritic structure. Phys. Rev. Lett. 78, 4665–4668 (1997)

26. Payne, T., Newmark, J., Reid, K.H.: The focally demyelinated rat fimbria: A new in vitro model for the study of acute demyelination in the central nervous system. Experimental neurology 114(1), 66–72 (1991)

27. Thaisetthawatkul, P., Logigian, E.L., Herrmann, D.N.: Dispersion of the distal compound muscle action potential as a diagnostic criterion for chronic inflammatory demyelinating polyneuropathy. Neurology 59(10), 1526–1532 (2002)

28. Isose, S., Kuwabara, S., Kokubun, N., Sato, Y., Mori, M., Shibuya, K., Sekiguchi, Y., Nasu, S., Fujimaki, Y., Noto, Y., et al.: Utility of the distal compound muscle action potential duration for diagnosis of demyelinating neuropathies. Journal of the Peripheral Nervous System 14(3), 151–158 (2009)

29. Rushton, W.: A theory of the effects of fibre size in medullated nerve. The Journal of physiology 115(1), 101–122 (1951)

30. London, M., Meunier, C., Segev, I.: Signal transfer in passive dendrites with nonuniform membrane conductance. Journal of Neuroscience (1999)

31. Koch, C.: Cable theory in neurons with active, linearized membranes. Biological Cybernetics 50, 15–33 (1984)

32. Stephanova, D.I., Kolev, B.D.: Computational Neuroscience: Simulated Demyelinating Neuropathies and Neuronopathies. CRC Press, ??? (2013)

33. Rall, W.: Membrane potential transients and membrane time constant of motoneurons. Experimental neurology 2(5), 503–532 (1960)

34. Tuckwell, H.C.: Introduction to Theoretic Neurobiology. Cambridge Univ. Press, Cambridge (1988)

35. Bostock, H., Sears, T.: Continuous conduction in demyelinated mammalian nerve fibres. Nature 263, 786–787 (1976)

36. Chow, C.C., White, J.: Spontaneous action potential fluctuations due to channel fluctuations. Bioph. J. 71, 3013–3021 (1996)

37. Faisal, A.A., White, J.A., Laughlin, S.B.: Ion-channel noise places limits on the miniaturization of the brain’s wiring. Curr Biol 15(12), 1143–9 (2005)

38. Katz, B., Schmitt, O.H.: Electric interaction between two adjacent nerve fibres. The Journal of physiology 97(4), 471–488 (1940)

39. Holt, G.R., Koch, C.: Electrical interactions via the extracellular potential near cell bodies. Journal of computational neuroscience 6(2), 169–184 (1999)

40. Binczak, S., Eilbeck, J., Scott, A.C.: Ephaptic coupling of myelinated nerve fibers. Physica D: Nonlinear Phenomena 148(1), 159–174 (2001)

41. Reutskiy, S., Rossoni, E., Tirozzi, B.: Conduction in bundles of demyelinated nerve fibers: computer simulation. Biological cybernetics 89(6), 439–448 (2003)

42. Chow, C.C., Kopell, N.: Dynamics of spiking neurons with electrical couplings. Neural Computation 12, 1643–1678 (2000)

43. van Kampen, N.G.: Stochastic Processes in Physics and Chemistry, 2nd edn. North-Holland, Amsterdam (1992)

44. Gerstner, W., Ritz, R., van Hemmen, J.: Why spikes? hebbian learning and retrieval of time-resolved excitation patterns. Biological cybernetics (1993)

45. Faisal, A., Laughlin, S.: Stochastic simulations on the reliability of action potential propagation in thin axons. PLoS Comput Biol 3, 79 (2007)

46. Plesser, H., Gerstner, W.: Noise in integrate-and-fire neurons: From stochastic input to escape rates. Neural Computation 12, 367–384 (2000)

47. Jolivet, R., Rauch, A., Lüscher, H., Gerstner, W.: Predicting spike timing of neocortical pyramidal neurons by simple threshold models. Journal of Computational Neuroscience 21, 35–49 (2006)

48. Gerstner, W., Naud, R.: How good are neuron models? Science 326, 379–380 (2009)

49. Bostock, H., Sears, T.: The internodal axon membrane: electrical excitability and continuous conduction in segmental demyelination. The Journal of Physiology 280(1), 273–301 (1978)

50. Kole, M.H., Letzkus, J.J., Stuart, G.J.: Axon initial segment kv1 channels control axonal action potential waveform and synaptic efficacy. Neuron 55(4), 633–647 (2007)

51. Rudolfer, S.M.: A markov chain model of extrabinomial variation. Biometrika 77(2), 255–264 (1990)

52. Lindén, H., Pettersen, K.H., Einevoll, G.T.: Intrinsic dendritic filtering gives low-pass power spectra of local field potentials. Journal of computational neuroscience 29(3), 423–444 (2010)

53. Einevoll, G.T., Kayser, C., Logothetis, N.K., Panzeri, S.: Modelling and analysis of local field potentials for studying the function of cortical circuits. Nat Rev Neurosci 14(11), 770–785 (2013)

54. Stephanova, D., Daskalova, M., Alexandrov, A.: Differences in potentials and excitability properties in simulated cases of demyelinating neuropathies. part i. Clinical neurophysiology 116(5), 1153–1158 (2005)

55. Stephanova, D., Daskalova, M.: Membrane property abnormalities in simulated cases of mild systematic and severe focal demyelinating neuropathies. European Biophysics Journal 37(2), 183–195 (2008)

56. Turrigiano, G.: Too many cooks? intrinsic and synaptic homeostatic mechanisms in cortical circuit refinement. Annual review of neuroscience 34, 89–103 (2011)

57. Seidl, A.H.: Regulation of conduction time along axons. Neuroscience 276, 126–134 (2014)

58. Waxman, S.G.: Axonal conduction and injury in multiple sclerosis: the role of sodium channels. Nature Reviews Neuroscience 7(12), 932 (2006)

59. Ritchie, J., Rang, H., Pellegrino, R.: Sodium and potassium channels in demyelinated and remyelinated mammalian nerve. Nature 294 (1981)

60. Rasband, M.N., Trimmer, J.S., Schwarz, T.L., Levinson, S.R., Ellisman, M.H., Schachner, M., Shrager, P.: Potassium channel distribution, clustering, and function in remyelinating rat axons. The Journal of Neuroscience 18(1), 36–47 (1998)

61. Platkiewicz, J., Brette, R.: A threshold equation for action potential initiation. PLoS computational biology 6(7), 1000850 (2010)

62. Coombes, S., Bressloff, P.: Saltatory waves in the spike-diffuse-spike model of active dendritic spines. Physical review letters 91(2), 028102 (2003)

63. Timofeeva, Y., Lord, G.J., Coombes, S.: Spatio-temporal filtering properties of a dendritic cable with active spines: a modeling study in the spike-diffuse-spike framework. Journal of Computational Neuroscience 21(3), 293–306 (2006)

64. Waxman, S., Brill, M., Geschwind, N., Sabin, T., Lettvin, J.: Probability of conduction deficit as related to fiber length in random-distribution models of peripheral neuropathies. Journal of the neurological sciences 29(1), 39–53 (1976)

65. Coggan, J.S., Prescott, S.A., Bartol, T.M., Sejnowski, T.J.: Imbalance of ionic conductances contributes to diverse symptoms of demyelination. Proceedings of the National Academy of Sciences 107(48), 20602–20609 (2010)

66. Halliday, A., McDonald, W., Mushin, J.: Delayed visual evoked response in optic neuritis. The Lancet 299(7758), 982–985 (1972)

67. Asselman, P., Chadwick, D., Marsden, D.: Visual evoked responses in the diagnosis and management of patients suspected of multiple sclerosis. Brain: a journal of neurology 98(2), 261–282 (1975)

68. Lachance, M., Longtin, A., Morris, C.E., Yu, N., Joós, B.: Stimulation-induced ectopicity and propagation windows in model damaged axons. Journal of Computational Neuroscience 3(37), 523–531 (2014)

69. Bégin, S., Bélanger, E., Laffray, S., Aubé, B., Chamma, É., Bélisle, J., Lacroix, S., De Koninck, Y., Côté, D.: Local assessment of myelin health in a multiple sclerosis mouse model using a 2d fourier transform approach. Biomedical optics express 4(10), 2003–2014 (2013)

